# Sensation is Dispensable for the Maturation of the Vestibulo-ocular Reflex

**DOI:** 10.1101/2024.05.17.594732

**Authors:** Paige Leary, Celine Bellegarda, Cheryl Quainoo, Dena Goldblatt, Başak Rosti, David Schoppik

## Abstract

Vertebrates stabilize gaze using a neural circuit that transforms sensed instability into compensatory counter-rotation of the eyes. Sensory feedback tunes this vestibulo-ocular reflex throughout life. Gaze stabilization matures progressively, either due to similar tuning, or to a slowly developing circuit component. Here we studied the functional development of vestibulo-ocular reflex circuit components in the larval zebrafish, with and without sensation. Blind fish stabilize gaze normally, and neural responses to body tilts mature before behavior. Instead, synapses between motor neurons and the eye muscles mature with a timecourse similar to behavioral maturation. Larvae without vestibular sensory experience, but whose neuromuscular junction was mature, had a strong vestibulo-ocular reflex. Development of the neuromuscular junction, and not sensory experience, determines the rate of maturation of an ancient behavior.

---

Early sensory deprivation profoundly disrupts neural function and associated behaviors in visual (*1*), auditory (*2*), and vestibular (*3, 4*) systems, and sensory feedback is responsible for tuning mature vestibular behaviors, (*5*). Sensory experience might therefore set the pace at which circuits and the behaviors they subserve emerge; alternatively, such functional maturation might reflect development of the underlying circuit components. While each component of a neural circuit plays a necessary role in generating behavior, a component’s necessity does does not confer rate-limiting status. Instead — by analogy to chemistry (*6*) — a rate-limiting component would be one whose development is the most protracted, determining the rate at which behavior can reach maturation. Despite recent technological improvements in connectomics for circuit identification (*7, 8*), the complexity and *in utero* development (*9*) of mammalian circuits complicates measuring rates of development, let alone linking them to behavior.

We studied an archetypal sensorimotor circuit that stabilizes gaze in a simple model vertebrate, the larval zebrafish. The relative simplicity and high conservation of this circuit across vertebrates (*10*) has made it a powerful model to uncover neural mechanisms of sensorimotor behavior (*11*). The vestibulo-ocular reflex circuit consists of sensory afferents, central interneurons, and motor neurons that together transform head/body tilts into counter-rotatory eye movements (*12*) (Figure 1A). When mature, this feed-forward circuit generates eye movements matching the head/body velocity, minimizing retinal slip and stabilizing gaze. In vertebrates, both gaze stabilization and vestibulo-ocular reflex circuit components mature during early development (*13*). Here we use the transparent, externally-developing larval zebrafish to determine the role of sensation in vestibulo-ocular reflex circuit development.

**Figure 1:**
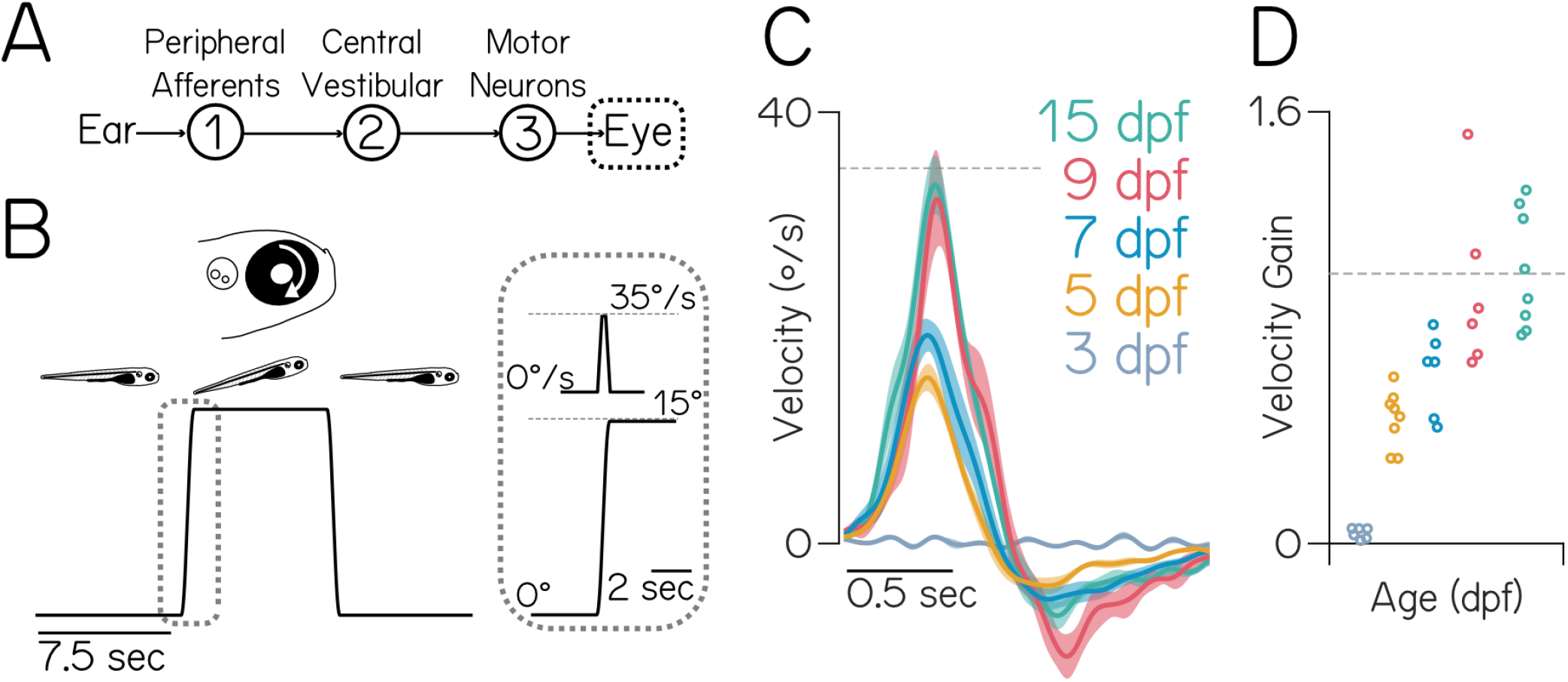
Gaze stabilization matures by 9 days post-fertilization (dpf). **(A)** Diagram of the feed-forward gaze stabilization circuit. Tilts are sensed by the utricle in the inner ear; information then flows via VIII^th^ cranial nerve peripheral afferents (1) to central vestibular neurons in the brainstem (2) to motor neurons in cranial nuclei nIII and nIV (3) that counter-rotate the eye. **(B)** Fish were rotated +15° (nose-up), held for 7.5 sec, then returned. The right eye rotates clockwise, or “down,” in response. Gray dotted box shows the trapezoidal velocity profile and angle of the tilt.**(C)** Average eye velocity ±SEM at 3, 5, 7, 9, and 15 dpf (N = 6, 8, 6, 6, 8 fish). Gray line at the peak tilt velocity (*±*35° /sec). **(D)** Gain (peak eye velocity / 35° /sec) for each fish, color indicates age as in C. Gains are significantly different between 3–5 dpf (*p*_*post-hoc*_ = 0.0021), 3–7 dpf (*p*_*post-hoc*_ = 6.51e-5), and 3–9 dpf (*p*_*post-hoc*_ = 4.57e-8), but not 9–15 dpf (*p*_*post-hoc*_ = 0.95). Horizontal line at gain = 1.

## Early maturation of gaze stabilization behavior

To determine when behavior matures, we measured the eyes’ response to body tilts across early development (Figure 1B). We previously used this approach to define the frequency response of the vestibulo-ocular reflex (*14*). Briefly, fish are immobilized with their eye freed and tilted in the pitch axis (nose-up/nose-down) on a rotating platform in complete darkness. Tilt magnitude (15°) and peak velocity (35°/sec) were selected to match the statistics of pitch tilts observed in freely swimming larvae (*15*). At and after 5 days post-fertilization (dpf), fish rotated their eyes down following nose-up pitch tilt stimuli (Figure 1C). A mature vestibulo-ocular reflex will counter-rotate the eyes at the same velocity as the body/head, eliminating residual image motion (retinal slip) and stabilizing gaze. We therefore parameterize behavioral performance as the ratio of the peak eye velocity to the peak platform velocity (gain). A gain of 1 is fully compensatory and mature, 0 indicates that the eye doesn’t move, and gains between 0–1 reflect immature and inadequate gaze stabilization. Gain improved with age from 3–9 dpf (*p*_*ANOVA*_ = 1.01e-9, Figure 1D), reached gains *∼*1 and did not change between 9–15 dpf (*p*_*post-hoc*_ = 0.95). These data indicate that behavior improves gradually over the first week of life, and performance plateaus *∼*9 dpf.

## Gaze stabilization develops normally in congenitally blind fish

As the vestibulo-ocular reflex minimizes retinal slip, we first asked how the absence of visual feedback during early life would impact the development of gaze stabilization behavior. Visual feedback is particularly important as fish ocular muscles do not contain proprioceptors (Golgi tendon organs and muscle spindles). We raised congenitally blind fish to *∼* 9 dpf (Methods) and measured eye rotations in response to body tilts, as in Figure 1 (Figure S1). Gain was comparable between blind fish and sighted siblings (*p*_*ANOVA*_ = 0.934). Despite its importance for calibrating gain in mature animals, visual information is therefore dispensable for the development of the vestibulo-ocular reflex.

## Vestibular neuron responses plateau well before behavioral performance peaks, with or without motor neuron-derived feedback

Since visual feedback is dispensable, either the slowest component of the circuit to develop and/or non-visual feedback could constrain behavioral maturation. Vestibular interneurons comprise the central node of a feed-forward circuit, where their activity encodes body/head tilt magnitude and direction (Figure 2A). To measure the development of vestibular responses to body tilts, we needed to record neural activity at eccentric body postures (Figure 2B), a challenge for conventional imaging with stationary microscopes. We used Tilt In Place Microscopy (TIPM)(*16*) to avoid rotating the microscope. Instead, TIPM returns fish from an eccentric orientation to the imaging plane faster (*∼* 5 msec) than the time constant of the calcium indicator (GCaMP6s)(*17*). The activity observed reflects the decay of the neuron’s eccentric response on return to horizontal. Central neurons in the tangential vestibular nucleus that project to extraocular motor nuclei nIII/nIV are indispensable for the vestibulo-ocular reflex (*14, 18*). They respond exclusively to either nose-up or nose-down tilts (*19*), and, when activated optogenetically, they induce torsional eye rotations (*18*). We performed longitudinal TIPM (*±* 25°) in these neurons to determine when their responses plateau. We found that vestibular neuron response strength increases dramatically between 3–5 (*p*_*t-test*_ = 1.2e-3, n=22 neurons/N=7 fish) but not between 5–7 or 7–9 dpf (*p*_*t-test*_ = 0.067, n=21/N=7, Figures 2C to 2E). While both behavior and vestibular neuron responses improve between 4–5 dpf, vestibular neuron response strength reaches a plateau well before behavioral performance is mature. The slowest component to mature must therefore be downstream of central vestibular interneurons.

**Figure 2:**
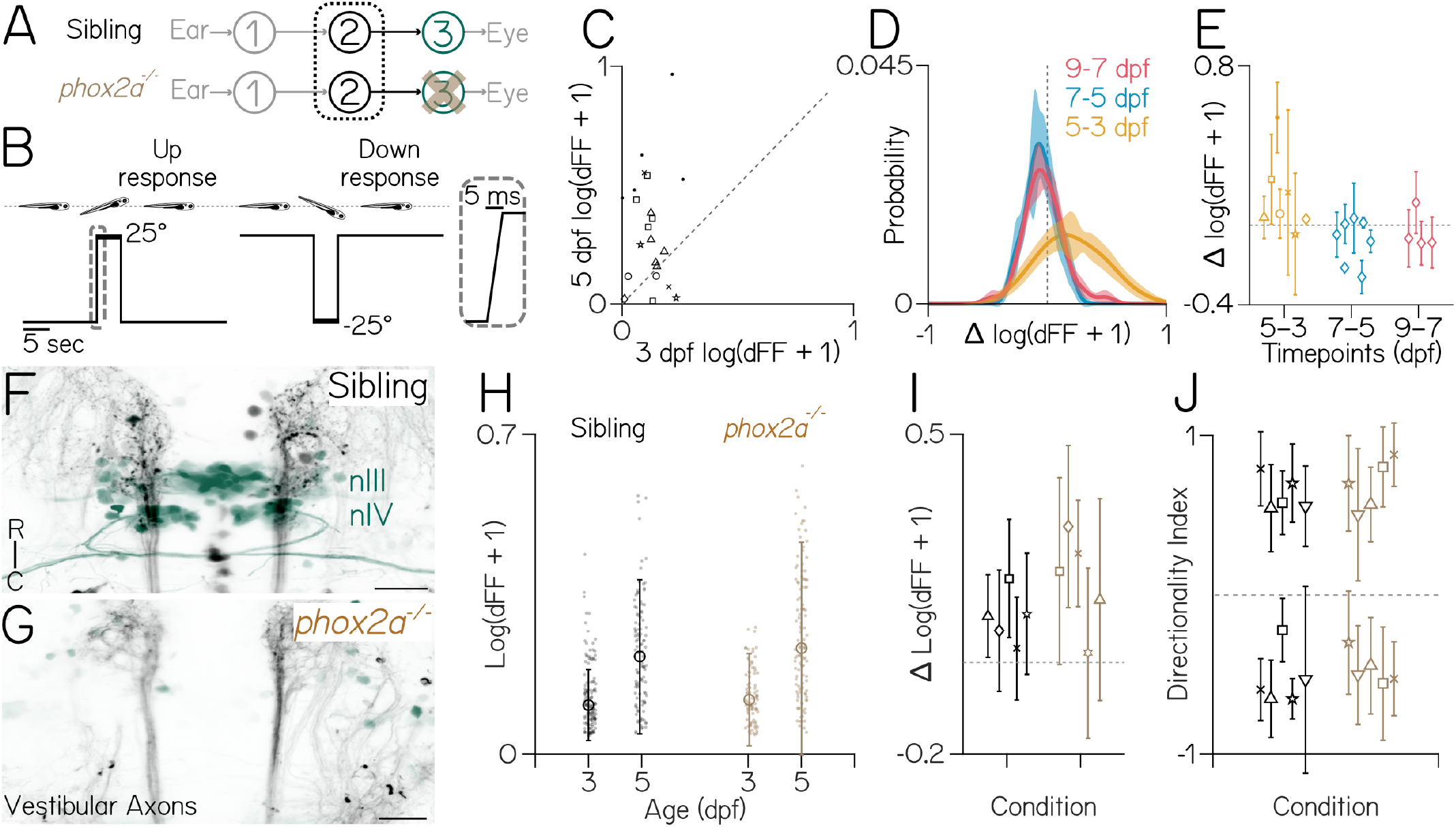
Central vestibular neuron responses plateau between 3–5 dpf, with or without motor neurons. **(A)** Diagram of the gaze stabilization circuit, focused on vestibular neurons in fish with (green) and without (brown, *phox2a*) motor neurons in cranial nuclei nIII and nIV. **(B)** A pitch-tilt stimulus trial with a 15 sec baseline, a rapid step (inset, 25° step), a 5 sec eccentric hold, and a rapid return for imaging. **(C)** Responses (dFF) to 25° nose-up steps are stronger at 5 dpf than 3 (*p*_*t-test*_ = 1.2e-3). Dashed line at 0. **(D)** Distribution *±*bootstrapped SD of pairwise differences of individual neurons between days (5-3, 7-5, 9-7 dpf) in response to 25° steps. **(E)** Data from Figure 2D, plotted as the median pairwise difference *±*IQR for neurons from each individual fish (N=7,7,4). 5–7 and 7–9 are not different from zero (*p*_*t-test*_ = 0.067 and 0.22). Dashed line at 0. **(F–G)** Axons of central vestibular nuclei (black) to motor neurons in nIII/nIV (green) in a 3 dpf sibling and *phox2a* mutant. Scale bar: 25µm. **(H)** Individual neuron responses (dots) to 19° pitch tilts in their preferred direction at 3 and 5 dpf. Median *±*IQR overlaid. Response changes between 3–5 dpf are significantly different for both siblings and mutants (*p*_*ANOVA*_ = 0), with no interaction between age and genotype (*p*_*ANOVA*_ = 0.053). **(I)** Data from Figure 2H, plotted as the median pairwise difference *±*IQR for paired neurons from each individual fish (n=113 neurons/N=5 siblings, n=116/N=5 *phox2a* mutants). Average response increased between 3–5 dpf with no effect of genotype on this increase (*p*_*ANOVA*_ = 5.03e-9 for age effect, *p*_*ANOVA*_ = 0.068 for interactions of age and genotype). Dashed line at 0. **(J)** Directionality indices of vestibular neurons at 5 dpf are comparable between *phox2a* null mutants (N=5 fish; n=102 neurons) and siblings (N=5,n=112), plotted as the median *±*IQR for each fish (*p*_*K-S test*_ = 0.66).

To determine whether motor-derived feedback influences the maturation of vestibular neuron responses, we adopted a loss-of-function approach. We performed TIPM to measure vestibular neuron responses on a *phox2a* mutant background (*20*) that fails to develop nIII/nIV extraocular motor neurons (Figures 2A, 2F and 2G). Following *±* 19° tilts, responses of individual vestibular neurons (Figure 2H) increased between 3–5 dpf for both sibling fish and mutants (*p*_*ANOVA*_ = 1.95e-26, age x genotype *p*_*ANOVA*_ = 0.053; 3 dpf: n=137 neurons over N=5 siblings, n=88/N=5 mutants; 5 dpf ; 5 dpf: n=113/N=5 siblings, n=116/N=5 mutants). Across fish (Figure 2I), the average response of paired (tracked) neurons increased between 3–5 dpf (*p*_*ANOVA*_= 0) with no effect of genotype (*p*_*ANOVA*_ = 0.069). Finally, to evaluate the development of peripheral vestibular inputs, we examined directional selectivity at 5 dpf. Consistent with a previous report (*19*), the ratio of nose-up and nose-down sensitive (directionality index values *<*|0.1|) neurons was nearly even, with no difference between genotype (*p*_*K-S test*_ = 0.66; 51.9 *±*6% nose-up, 48 *±*6% nose-down, n=118 neurons over N=5 siblings; 48.1*±*13% nose-up stimuli, 51.8±13% nose-down, n=117/N=5 *phox2a* mutants) (Figure 2J). These results establish that, even without motor neurons — and by extension, motor-derived feedback — vestibular neuron responses develop early relative to behavior.

## Extraocular motor neuron responses develop between 3–5 dpf

As the last central node in a feed-forward reflex, extraocular motor neuron activity reflects all sensory inputs from the entire circuit (Figure 3A). Consequentially, if the development of motor neuron responses reaches a plateau earlier than behavioral improvement, the slowest developmental process must be in the motor periphery. We repeated longitudinal TIPM (*±* 25°) to determine when motor neuron responses plateau. However, as the calcium responses of motor neurons have not been evaluated in response to body tilts, we first characterized the tuning and sensitivity properties of these neurons to a gradient of stimulus steps (Figure 3B). To match the pulling direction of the muscle, superior oblique motor neurons should respond predominantly to nose-up tilts. As expected, superior oblique neurons at 5 dpf responded unidirectionally to nose-up body tilts (Figure 3C) and not to nose-down body tilts (Figure 3D). Motor neuron activity at 5 dpf varied as a function of nose-up tilt eccentricity (n=22/32 neurons with slope *>* 0, *p*_*post-hoc*_ = 0.002, Figure 3E).

**Figure 3:**
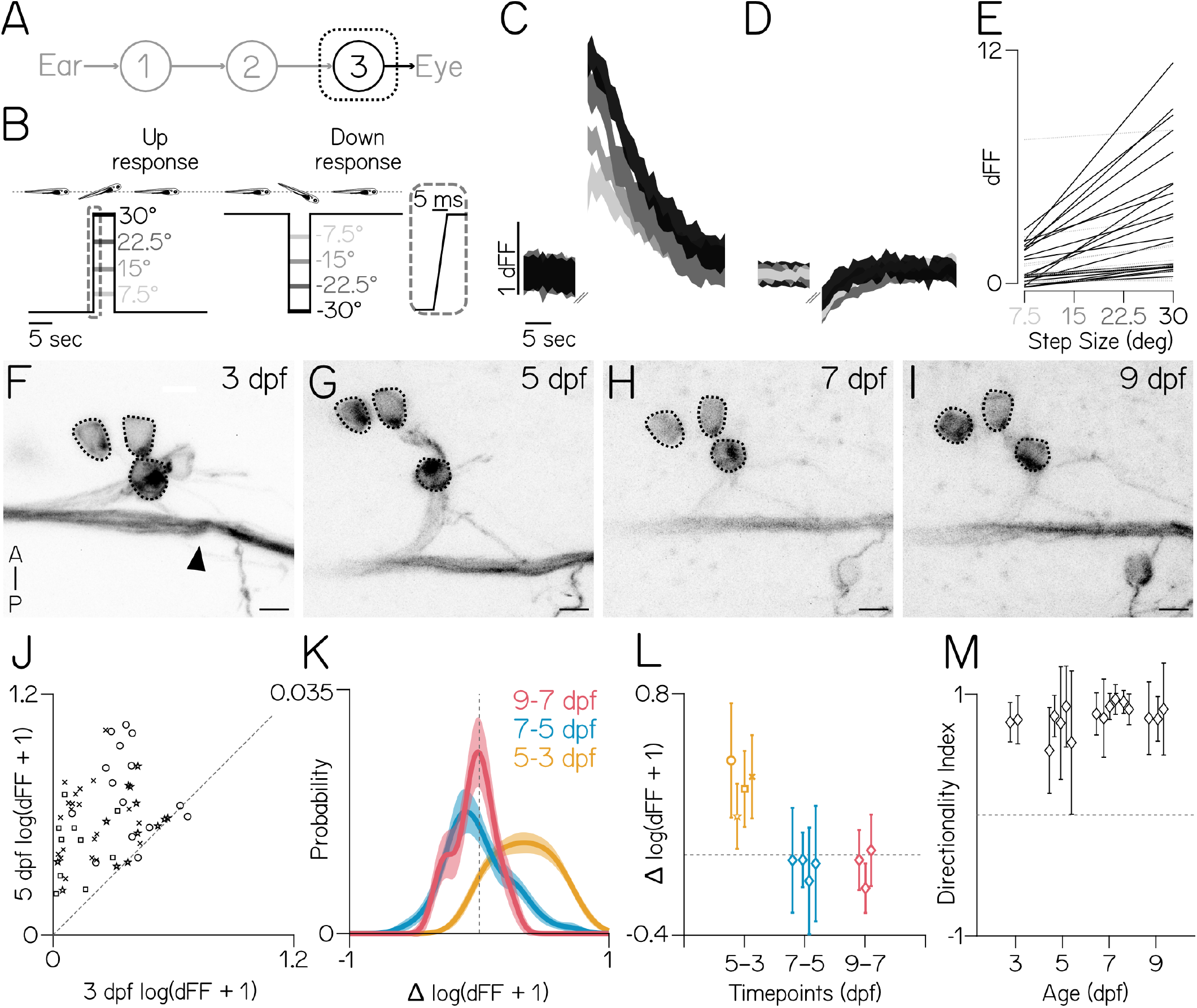
Superior oblique motor neuron responses plateau between 3–5 dpf. **(A)** Diagram of the gaze stabilization circuit, focused on motor neurons. **(B)** A pitch-tilt stimulus trial with a 15 sec baseline, a rapid step (inset, 30° step), a 5 sec eccentric hold, and a rapid return for imaging. **(C–D)** Mean *±*SD of responses (dFF) to nose-up (C) and nose-down (D) pitch steps from a single superior oblique extraocular motor neuron at 5 dpf (3–5 trials per step size). Pitch amplitude (grayscale) as in Figure 3B. **(E)** Best-fit lines of responses (dFF) from superior oblique motor neurons to nose-up pitch tilts. Black lines have slopes *>*0 (n=22/32, N=3 fish). **(F–I)** Three superior oblique extraocular motor neurons (dotted circles) and the trochlear nerve (black arrow) tracked from 3–9 dpf. Scale bar = 5µm. **(J)** Responses (dFF) to 25° nose-up steps are greater at 5 dpf than 3 dpf (*p*_*t-test*_ = 1.31e-16). Dashed line at 0. **(K)** Distribution *±*bootstrapped SD of pairwise differences of individual neurons between days (5-3, 7-5, 9-7 dpf) in response to 25° nose-up steps. **(L)** Data from Figure 3K, plotted as the median pairwise difference for neurons from each individual fish (N=4,4,3). 5–7 and 7–9 are not different from zero (*p*_*t-test*_ = 0.69, 0.23). Dashed line at 0. **(M)** Median *±* IQR directionality index to *±*25° steps do not change with age (*p*_*K-W*_ = 0.09). +1 is selective for nose-up, -1 nose-down. Dashed line at 0.

To measure how tilt responses developed, we recorded calcium signals from a transgenic line with sparse expression in superior oblique motor neurons (Figure S2). Motor neuron somata were stably positioned within nIV (Figures 3F to 3I), allowing reliable identification of the same neurons over two-day increments: 3–5 dpf, 5–7 dpf, and 7–9 dpf. Almost every superior oblique neuron had a stronger response to 25° nose-up body tilts at 5 dpf compared to 3 (*p*_*t-test*_ = 1.31e-16, n=59 pairs over N=4 fish, Figure 3J). While the distributions of differences varied between 5–7 and 7–9 (*p*_*K-S*_ = 5.7e-15 Figure 3K), when evaluated across fish, the average response did not change between 5–7 (*p*_*t-test*_ 0.69, n=53/N=4) or 7–9 (*p*_*t-test*_ = 0.23, n=23/N=3, Figure 3L). We gathered another dataset with 19° nose-up and nose-down steps to evaluate the development of directional selectivity (Methods). We saw no changes across time (*p*_*K-W*_ = 0.09, 3 dpf: n=16 neurons over N=2 fish; 5 dpf: n=32/N=5; 7 dpf: n=60/N=7; 9 dpf: n=30/N=3, Figure 3M). Notably, all of the brain’s activity related to the vestibulo-ocular reflex (*21*) must converge on extraocular motor neurons. Since extraocular motor neuron response strength and directional selectivity appear to plateau by 5 dpf — well before behavior — we conclude that the slowest component of the circuit to develop must lie downstream of motor neurons.

## The developmental timecourse of the postsynaptic neuromuscular junction matches the maturation of behavior

We next focused on the extraocular neuromuscular junction (Figure 4A). We labeled postsynaptic acetyl-choline receptors with fluorescently-conjugated alpha-bungarotoxin (*α*-BTX, Figures 4B to 4D) and presy-naptic vesicles with an SV2A antibody (*22*) (Figures 4E to 4G). We targeted all four eye muscles used for vertical/torsional gaze stabilization: superior oblique (SO), superior rectus (SR), inferior oblique (IO), inferior rectus (IR) in 3–9 and 14 dpf fish. Over time, the fraction of each muscle labelled with *α*-BTX increased with a timecourse comparable to behavioral improvement (p_ANOVA_ = 0) (SO, n=106 muscles over N=61 fish; SR n=52/N=32; IR n=43/N=28; IO n=38/N=26 Figure 4H), with no difference in labeling observed between 9 and 14 dpf (*p*_*post-hoc*_ = 0.185). In contrast, SV2A labeling appeared earlier and developed more rapidly. SV2A labeling significantly increased between 3–5 dpf (*p*_*post-hoc*_ = 5.5e-4), but did not change afterwards: (*p*_*post-hoc*_ *>* 0.13 for 5–7,5–9,5–14,7–9,7–14,9–14 SO, n=34/N=17; SR n=33/N=17; IR n=29/N=17; IO n=28/N=15 Figure 4I).

**Figure 4:**
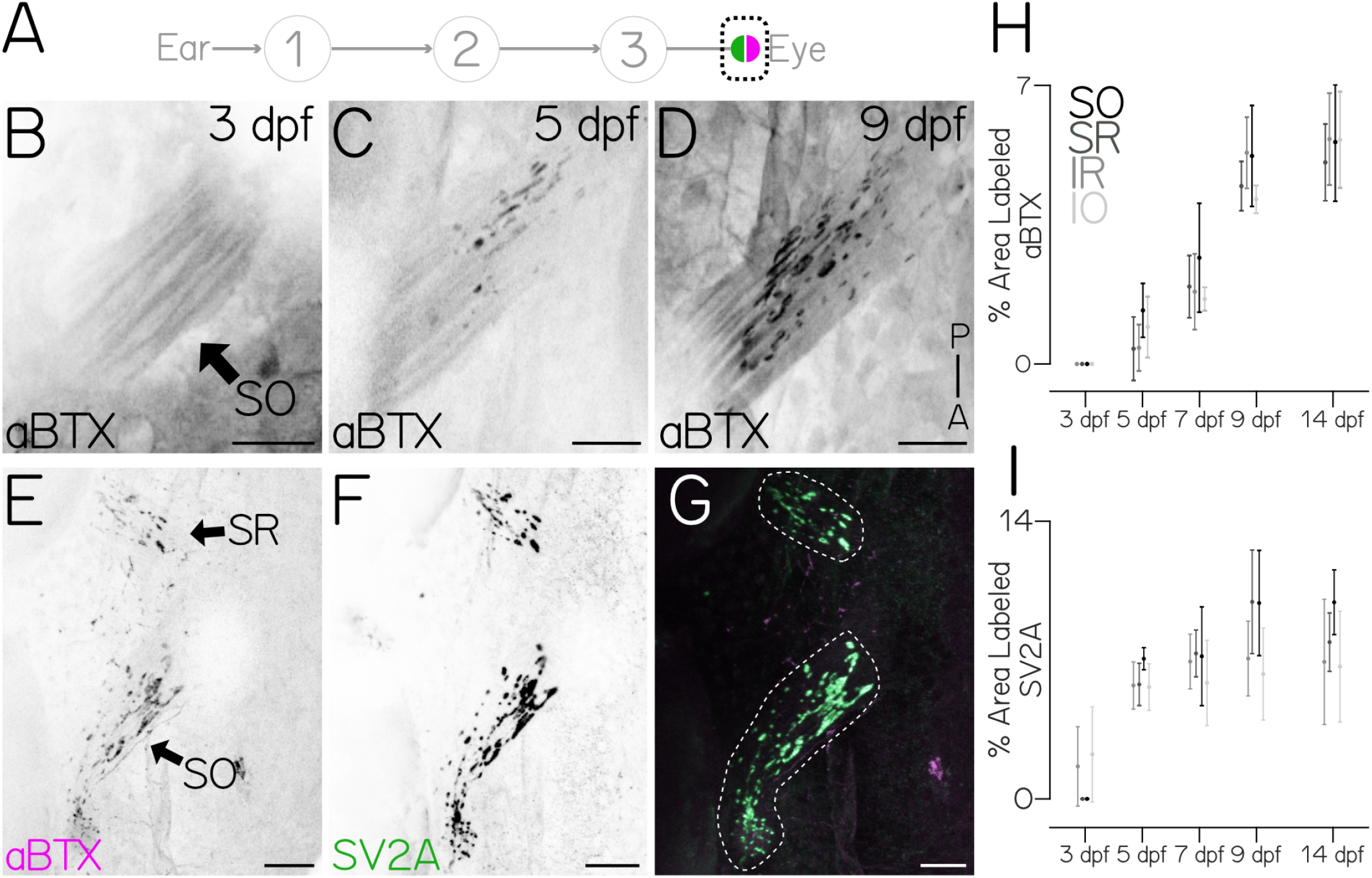
The timecourse of post-synaptic neuromuscular junction development matches behavioral maturation. **(A)** Diagram of the gaze stabilization circuit, focusing on pre-(green) and postsynaptic (magenta) components of the neuromuscular junction. **(B-D)** Dorsal projection of the superior oblique muscle (black arrow) at 3 (B) 5 (C) and 9 (D) dpf labeled with fluorescent (*α*-BTX). Scale bar = 20µm **(E-G)** Comparison of superior oblique (SO, arrow) and superior rectus (SR, arrow) pre-(SV2A, green) and postsynaptic (*α*-BTX, magenta) label at 8 dpf. Dotted line in (G) denotes muscle bound used for quantification. SO muscle: 6.59% *α*-BTX and 15.03% SV2A. SR muscle: 4.98% *α*-BTX and 9.33% SV2A. Scale bar = 20µm. **(H)** Postsynaptic *α*-BTX label in SO, SR, inferior rectus (IR) and inferior oblique (IO) muscles. Dots are the mean *±*SD area labeled. For all muscle types, labeling increased between 3 and 9 dpf (*p*_*ANOVA*_ = 0), but not between 9 and 14 dpf (*p*_*post-hoc*_ = 0.185). **(I)** Presynaptic SV2A staining in SO,SR,IR,IO. Dots are the mean *±*SD area labeled. For all muscle types, labeling significantly increased between 3–5 dpf (*p*_*post-hoc*_ = 5.5e-4), but not following 5 dpf (5–7 *p*_*post-hoc*_ = 0.889, 7–9 *p*_*post-hoc*_ = 0.462, 9–14 *p*_*post-hoc*_ = 0.983).

Larval zebrafish eye muscles assume their adult configuration by 3 dpf, with both thick and thin myofibrils by 4 dpf (*23*), allowing the eye to assume static orientations up to *±*30° (*14*), and a mature optokinetic response (*<*0.5Hz) by 6 dpf (*24*). To evaluate whether the eyes’ velocity was constrained by the properties of the muscle, we measured eye movements at 5 dpf following unnaturally rapid tilts (90°/sec, 600°/sec^2^ instead of 35°/sec, 150°/sec^2^, Figure S3A). We found that although gaze stabilization behavior is far from mature, and the *α*-BTX signal has just begun to emerge, the eye could rotate faster (*p*_*t-test*_ = 0.016, Figures S3B and S3C). Together these findings implicate post-synaptic development at the neuromuscular junction as the slowest step in circuit maturation.

## Vestibulo-ocular reflex behavior is mature following restoration of transient sensory deprivation

If sensory experience sets the rate of behavioral development, then transient loss of vestibular sensation should delay normal improvements to gaze stabilization. On restoration, vestibulo-ocular reflex performance should steadily improve. Alternatively, if behavioral performance reflects the capacity of the circuit component that matures most slowly, then on restoration of vestibular sensation, behavior would be immediately comparable between experienced and transiently-deprived larvae.

We investigated the emergence of the vestibulo-ocular reflex in a mutant line (*otogelin*) that, in some cases, is transiently insensitive to body tilts. Under normal conditions (*19*), larval zebrafish rely exclusively on the utricle to sense body tilts (*25*), which uses an inertial difference between an otolith and hair cells embedded in a gelatinous macula to transduce linear acceleration. Initial calcification of the utricular otolith normally occurs between 18-24 hours post-fertilization (hpf) (*26*). The majority of *otogelin* mutants do not generate a utricular otolith and, in the dark, are tilt-blind (*27*). A small fraction of *otogelin* mutants show delayed generation of the utricular otolith, which appears over a 24 hr period around two weeks of age (Figures 5B and 5C), at which point postural behaviors resume (*28*). We hypothesized that since the neuromuscular junction would have time to mature, *otogelin* mutants should show comparable behavior to siblings as soon as the utricle becomes functional.

**Figure 5:**
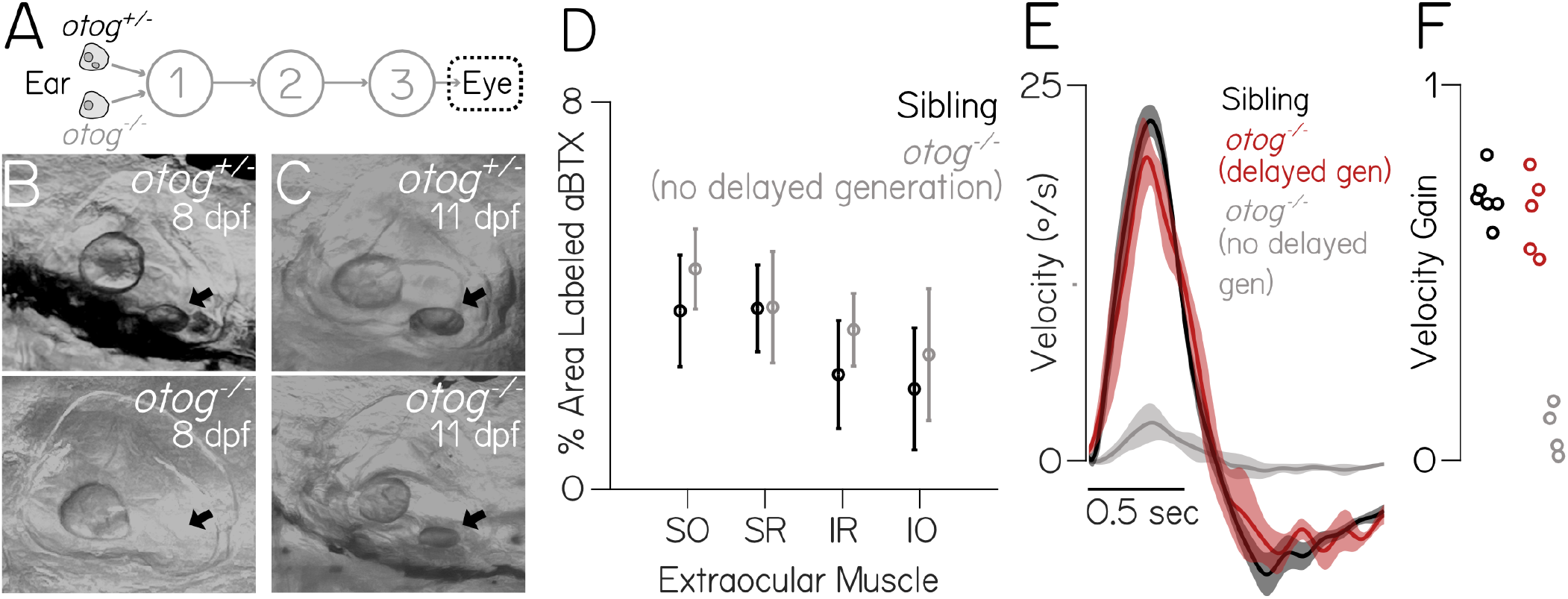
Gaze stabilization behavior emerges rapidly after delayed emergence of tilt sensation. **(A)** Diagram of the gaze stabilization circuit illustrating *otogelin*^*-/-*^ phenotype with focus on eye movements. **(B-C)** Top: *otogelin*^+/-^, utricular otolith visible in ear at 8 (left) and 11 dpf (right, black arrows). Bottom: *otogelin*^-/-^ fish without a utricular otolith at 8, and with at 11 dpf (black arrows). **(D)** Postsynaptic *α*-BTX label from sibling and *otogelin*^*-/-*^ fish in SO (N=6,6), SR (N=5,6), IR (N=5,5), IO (N=4,6). **(E)** Average eye velocity traces*±* SEM to a nose-up step as in Figure 1B. **(F)** Vestibulo-ocular reflex gain (peak eye / 35°/sec) for each fish (N=6,5,4) in Figure 5E. Gains did not differ between siblings and regenerated *otogelin*^*-/-*^ (*p*_*post-hoc*_ = 0.66), unlike *otogelin*^*-/-*^ (*p*_*post-hoc*_ = 1.84e-7 and 6.20e-7).

We first confirmed that the neuromuscular junction developed normally in *otogelin* mutants; we observed strong *α*-BTX labeling at 8 dpf in both siblings and mutants in all four muscles (Figure 5D). Next, we raised and screened *>*2000 *otogelin* mutants daily to identify 5 fish with nascent bilateral utricular otoliths between 11–16 dpf (Methods). Eye movements were measured following body tilts (Figure 1B) on the day after identification, when the otolith had reached normal size. Performance was statistically indistinguishable between siblings (N=6) and mutants (N=5) with newly generated otoliths (*p*_*post-hoc*_ = 0.66, Figures 5E and 5F). In contrast, 11–16 dpf fish that never developed otoliths were profoundly impaired (*p*_*post-hoc*_ = 1.84e-7 sibling vs. no otolith, 6.20e-7 delayed otolith vs. no otolith). We infer from the rapid emergence of functional gaze stabilization in *otogelin* mutants that initial vestibular circuit assembly can proceed without sensory input, consistent with other reports (*29, 28, 30*). Our data show that vestibular experience is dispensable for maturation of the vestibulo-ocular reflex.

## DISCUSSION

We find that, remarkably, both the vestibulo-ocular reflex circuit and behavior can mature without any vestibular or visual input at all. This finding is particularly striking given that gaze stabilization is plastic later in life: visual feedback following eye movements is used to adjust the sensitivity of central vestibular neurons to modulate gain (*5*). Even when there are no motor neurons to move the eyes, we see that central vestibular neuron responses to tilts are unchanged, underscoring the early dispensability of sensory feedback. In the light, the vestibulo-ocular and optokinetic reflexes work together to stabilize gaze; feedback following optokinetic eye movements might suffice to shape the vestibulo-ocular reflex circuit we studied here. However, such modulation would be inconsistent with our finding that the vestibulo-ocular reflex matures normally in blind fish. Studies of locomotor development over the last century underscore the importance of feedback as animals learn to move properly in their environment (*31, 32*). Undoubtedly, given the considerable morphological and neurological changes that happen between larval and adult stages, feedback will be similarly important to maintain an excellent vestibulo-ocular reflex. We show that that such plasticity acts on a mature scaffold that can evoke excellent behavior as soon as the neuromuscular junction is sufficiently developed.

By definition, evolutionary adaptations that support earlier/later development of sensorimotor reflex behaviors must manifest with changes at the circuit component that is slowest to develop (i.e. rate-limiting). Does the rate of development of the neuromuscular junction limit maturation of other behaviors? Early neuromuscular junction development permits developing rodents to make spontaneous limb contractions, or “twitches,” that can shape sensory cortical (*33*) and cerebellar development (*34*). Similar twitches (*35*) precede development of the hindbrain neurons responsible for evoked escape swimming in larval zebrafish (*36*), the earliest sensorimotor reflex behavior; by inference, the rate-limiting process for escapes is upstream of the motor periphery. Differences in the location of the rate-limiting process may reflect selective pressures that differ between precocial (mobile at birth) and altricial (immobile) animals. Empirically, when the pressure is on to stand/run/swim or be eaten, the motor periphery develops much more rapidly to ensure that “the neuro-muscular system [is] ready for use when the brain needs it” (*37*). We suggest that defining the location of rate-limiting processes will speak to how circuits came to be.

Recent advances in transcriptomics and connectomics (*7, 8*) revealed the elements of and interactions between vertebrate neural circuit nodes. Similar advances transformed longitudinal (*38*) and circuit-level (*39*) measurements linking neuronal activity and behavior (*40*). In contrast, conceptual frameworks (*30*) lag behind, relying primarily on classic loss-of-function approaches to establish necessity (*41*). Here, motivated by a powerful simplification in chemistry (*6*), we propose a candidate rate-limiting process for the maturation of a vertebrate sensorimotor behavior. Our approach moves beyond “necessity,” linking neural circuit development to behavioral performance.

## ACKNOWLEDGMENTS

The authors would like to thank Cecilia Moens and Martha Bagnall for all the fish, Hannah Gelnaw for assistance with husbandry, and Katherine Nagel, Jeremy Dasen, Niels Ringstad, Lynne Kiorpes, Michael Long, György Buzsáki, Alexander Schier, Claude Desplan, Steve Burden along with the members of the Schoppik and Nagel labs for their valuable feedback and discussions.

## AUTHOR CONTRIBUTIONS

Conceptualization: PL and DS, Methodology: PL, CB, CQ, DG, BR, and DS, Investigation: PL, Visualization: PL, Writing: PL and DS, Editing: PL and DS, Funding Acquisition: PL and DS, Supervision: DS.

## Data Availability

All raw data and analysis code are available at the Open Science Foundation, DOI: 10.17605/OSF.IO/7Z5UP

## Supplementary Materials

Materials and Methods

Figs. S1 to S3

**Figure S1:**
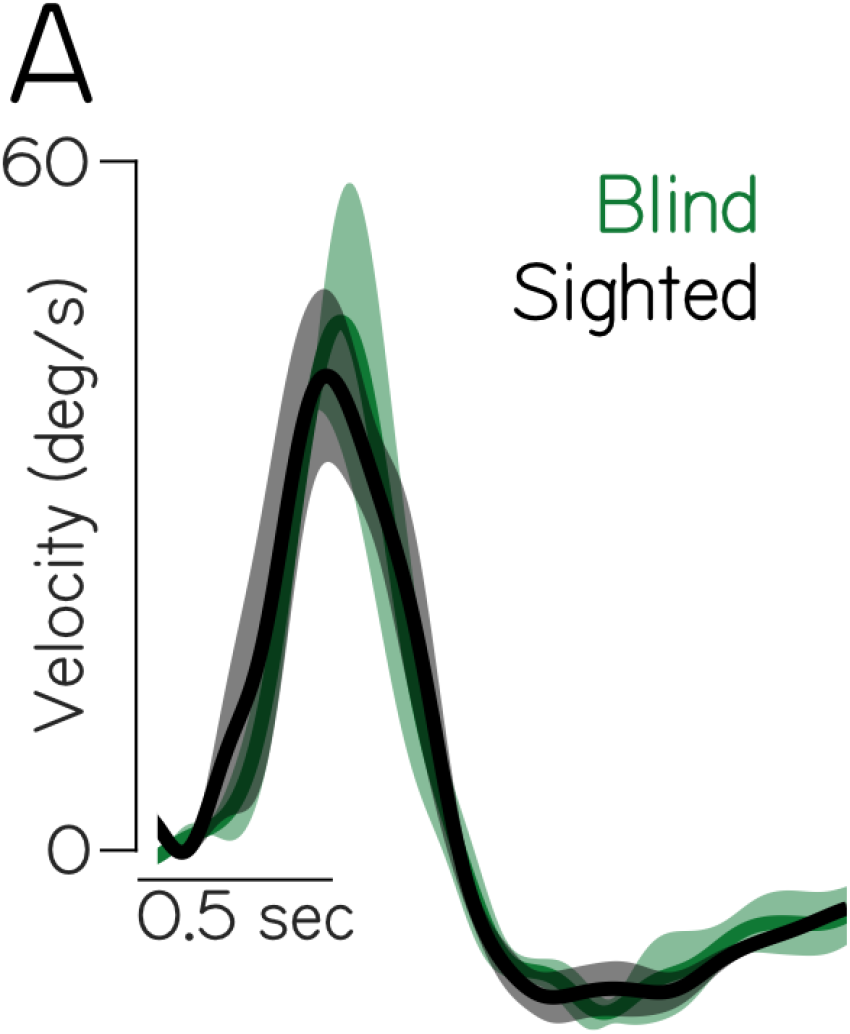
Congenitally blind fish achieve comparable gaze stabilization to sighted siblings. **(A)** Eye velocity traces*±*SEM of six fish morphologically aged approximately 9 dpf stepped to +15°at peak velocity 35°/sec. Congenitally blind fish traces in green (N = 3), sighted siblings in black (N = 3).

**Figure S2:**
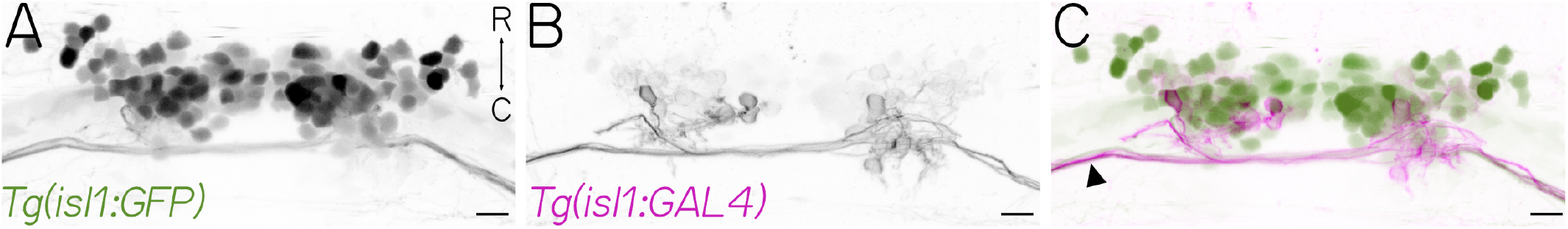
Sparse labeling of superior oblique extraocular motor neurons. **(A-C)** Dorso-ventral maximum intensity projection of the extraocular motor neurons of cranial nuclei nIII and nIV visualized in a 4 dpf fish using the triple transgenic line *Tg(isl1:GFP)* (A, green) crossed to *Tg(isl1:GAL4)*;*Tg(UAS:KillerRed)* (B, magenta). Scale bar = 10µm.

**Figure S3:**
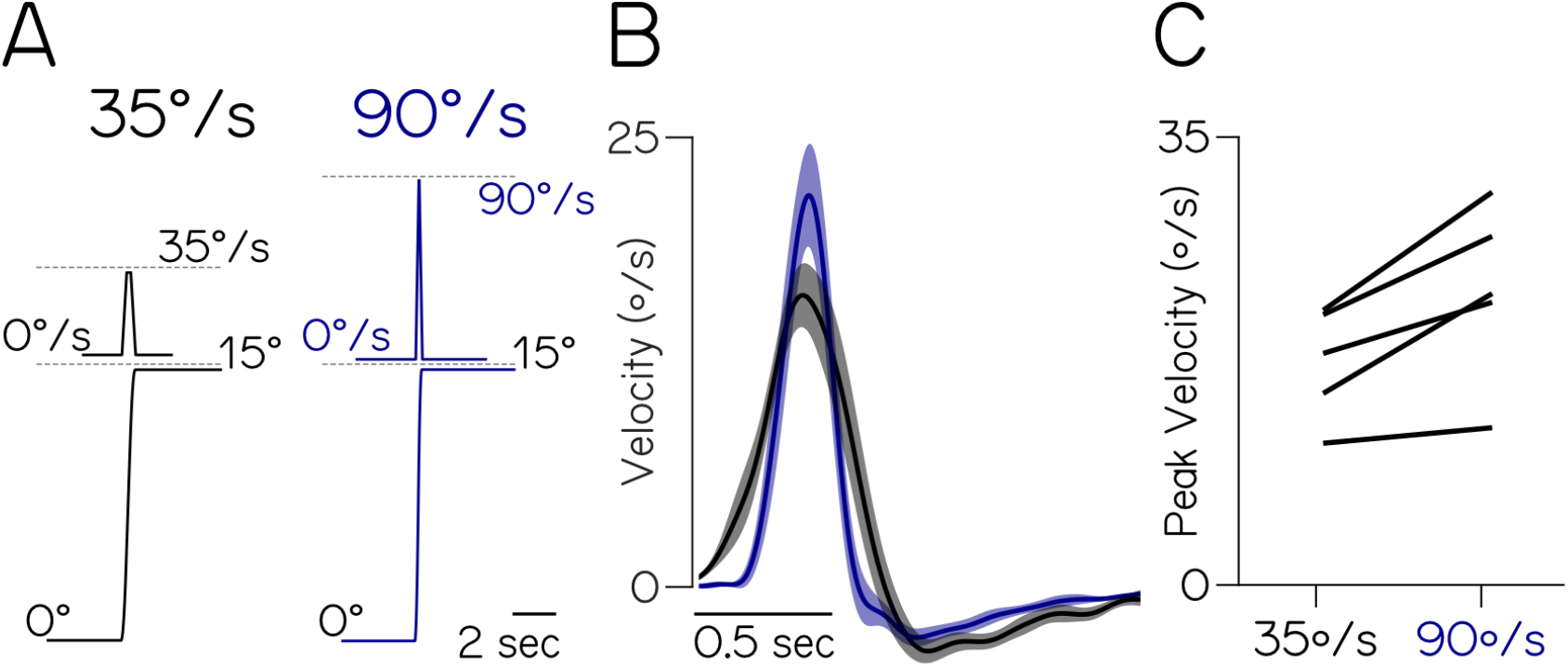
Stronger tilt stimulation evokes faster eye rotations before gaze stabilization behavior is fully mature. **(A)** Schematic of our standard (left, black) table-tilt stimulus and a faster (right, blue) variant, as described in Figure 1A. Fish are stepped upwards to a a +15° (nose-up) step, held for 7.5 sec, then returned. In black, the peak velocity is 35°/sec and the peak acceleration is 150°/sec. In blue, the peak velocity is 90°/sec and the peak acceleration is 600°/sec. Traces derived from generated stimulus steps. **(B)** Eye velocity traces *±*SEM of five fish at 5 dpf stepped using normal (black) or fast (blue) stimuli. **(C)** Mean peak eye velocity of 5 dpf fish for normal (black) and fast (blue) tilts. Peak eye velocities are significantly larger after fast tilts (*p*_*t-test*_ = 0.016, N=5).

## MATERIALS AND METHODS

### Fish Care

All procedures involving zebrafish larvae (*Danio rerio*) were approved by the New York University Langone School of Medicine Institutional Animal Care and Use Committee (IACUC). Fertilized eggs were collected and maintained at 28.5°C on a standard 14h light/10 h dark cycle. Before 5 dpf, larvae were maintained at densities of 20–50 larvae/10-cm-diameter Petri dish, filled with 25–40 mL of E3 with 0.5 ppm methylene blue. After 5 dpf, larvae were maintained at densities under 20 larvae/Petri dish and were fed cultured rotifers (Reed Mariculture) daily. Rotifer density for blind larvae was increased *∼*50-fold to facilitate feeding. Consequentially, blind fish ate more and developed more rapidly such that body length at 6 dpf was comparable to 9 dpf wild-type larvae.

### Transgenic lines

Functional imaging and confocal experiments were conducted on the *mitfa*^-/-^ background to remove pigment. Two-photon calcium imaging experiments were performed on the *Tg(isl1:GAL4)*^*fh452*^;*Tg(14xUAS:GCaMP6s)* background (*42, 43*). Confocal imaging experiments used larvae of Tg(isl1:GFP) background (*44*). *α*-bungarotoxin and SV2A labeling experiments used larvae from the *roy* (*mpv17*^a9/a9^) background (*45*) to eliminate autofluorescence from iridiphores of the eye. Vestibular projection neuron functional imaging was performed using the *Tg(−6*.*7Tru*.*Hctr2:GAL4-VP16)* line (*46*). All larvae used were selected for brightness of fluorescence relative to siblings to facilitate imaging experiments. Mendelian ratios were observed among selected/unselected larvae; we infer that selected larvae were homozygous for a given allele. Motor neuron loss-of-function experiments were performed on larvae from F3 or older generations of *phox2a* mutants (*20*), screened on the *Tg(isl1:GFP)* background for the presence or absence of fluorescence in nIII/nIV. *oto-gelin* mutants were rks^vo66/vo66^(*27*). Blind fish were from the *lakritz* (*atoh7*^*th241/241*^) background that does not develop retinal ganglion cells (*47*). *Tg(14xUAS-E1B:KillerRed)* (*48*) was used for visualization.

### Confocal Imaging

Larvae were anesthetized in 0.2 mg/mL ethyl-3-aminobenzoic acid ethyl ester (MESAB, Sigma-Aldrich E10521, St. Louis, MO) prior to imaging. Larvae were mounted dorsal side-up (axial view) in 2% low-melting point agarose (Thermo Fisher Scientific 16520) in E3. Images were collected on a Zeiss LSM800 confocal microscope with a 20× water-immersion objective (Zeiss W Plan-Apochromat 20x/1.0). Anatomical stacks of nIII/nIV spanned approximately 100µm in depth, sampled every two microns. Image stacks were analyzed using Fiji/ImageJ (*49*).

### Tilt In Place Microscopy (TIPM)

TIPM is an imaging technique to measure neural responses to body tilts by rapidly rotating larvae (peak angular velocity, *<*3,000°/s) and peak acceleration (*>*1e7°/s^2^) (*16*). Briefly, larvae aged 3, 5, 7, or 9 dpf were mounted dorsal-up in 2% low-melt agarose in E3 in the center of the mirror of a large beam diameter single-axis scanning galvanometer system (ThorLabs GVS011, 10 mm 1D, 40° range). A single trial consisted of of a 15 sec pre-stimulus baseline, a rapid (*∼*5 msec) eccentric step, 5 sec at the eccentric tilt, a rapid (*∼*5 msec) return to baseline, and a 20 sec period at the horizontal. Experiments mapping the sensitivity of motor neurons consisted of 4–5 repeats of 7.5°, 15°, 22.5°, or 30° steps, order randomized, lasting *∼*2 hrs/fish. To use the full range of the galvonometer, sensitivity-mapping stimuli were presented in a single pitch direction (nose-up or nose-down) at a time; direction order was randomized. Longitudinal experiments were performed similarly, using a single 25° step presented 4–5 times, lasting *∼*1 hr/fish. All experiments took place in complete darkness.

All stimuli were presented while imaging with a two-photon microscope (ThorLabs Bergamo) using a 20x water immersion objective (Olympus XLUMPLFLN20xW 20x/1.0). Illumination consisted of a pulsed infrared laser (Spectra-Physics MaiTai HP) at 920 nm using 6.1–18.8 mW of power, measured at the sample using a power meter (ThorLabs S130C).

For motor neuron imaging, volumes were centered over nIV in both hemisphere, with the trochlear nerve visible in each stack. Each volume consisted of 4–6 planes spanning 70µm in the dorso-ventral axis. Because expression was sparse and variegated in the *Tg(isl1:GAL4)*; *Tg(UAS:GCaMP6s* line, the imaging window was optimized to cover all labelled nIV cells for each fish. Imaging was performed at *∼*3 volumes/sec (*∼*26 frames/sec, *∼*74–150 x 50µm frame size, and 1µsec pixel dwell time).

For imaging of vestibular neurons in the tangential nucleus in *phox2a*^-/-^ fish and siblings, experiments were performed at 3 and 5 dpf. Larvae were sorted at 2 dpf into *phox2a*^-/-^ and sibling controls (non-phenotypic) based on the absence or presence of labeled extraocular motor neurons using the *isl1:GFP* transgene. Imaging volumes were centered over the tangential nucleus (*19*), one hemisphere at a time. Each volume consisted of 4-5 planes spanning 30µm in the dorso-ventral axis. Imaging was performed at *∼*3 volumes/second (*∼*20frames/second, *∼*100×60µm frame size, and 1µs pixel dwell time).

### Analysis of TIPM data

Volumes were pre-processed using Fiji/ImageJ (*49*). Each volume was motion-corrected to account for any slight shifts in the X or Y axes across multiple trials(*50*). Polygonal regions of interest (ROI) were drawn manually around all neuron somata visible in an average intensity projection. All neurons were included for analysis.

All subsequent analyses were performed using Matlab R2020b (MathWorks, Natick, MA, USA). Raw fluorescence traces from ROIs were extracted and, when imaging zoom had varied, normalized by ROI size to compensate for different pixel sizes. A neuron’s response was defined as the normalized change in fluorescence (dFF). dFF was calculated by taking the peak during the first second on return to horizontal, subtracting the baseline, and dividing by the baseline. Baseline fluoresence was defined as the average value over the first 10 frames at the beginning of each trial. To quantify the strength of directional tuning, a directionality index was calculated for each neuron. The directionality index (from -1 to 1) is defined as the difference in up/down dFF normalized by their sum. A neuron was defined as “tuned” if the absolute value of the directionality index was *>*0.1. Data are presented as log(dFF+1) to better visualize the full range of responses.

### Tracking Individual Neurons Across Time

Motor neurons and tangential nucleus neurons were manually tracked across three time points: 3–5 dpf, 5–7 dpf, and 7–9 dpf. TIPM data was used to define the soma position and, for motor neurons, the distance from the trochlear nerve. To ensure reliability, neurons across time were tracked independently by two separate experimenters (P.L. and either C.B. or C.Q.); only neurons where experimenters agreed were included. After 5 dpf, somata remained stationary, facilitating tracking. The majority of neurons recorded following 5 dpf were identifiable at the second time points (64.4 *±*19.4% between 5 and 7 dpf, 53.7*±*15.2% between 7 and 9 dpf). Before 5 dpf, some soma shifting was observed (typically in a caudal-to-rostral direction for motor neurons), and therefore an additional anatomy stack was taken at 4 dpf to more facilitate tracking neurons across time. The yield of confidently tracked neurons between 3 and 5 dpf was 59.7*±*19.9%.

### Whole-Mount Immunohistochemistry and Synaptic Labeling

Larvae were fixed and permeabilized as described in (*51*). Briefly, fish were anesthetized in 0.2 mg/mL MESAB then fixed in 4% paraformaldehyde overnight at 4°C. Following overnight fixation, fish were washed in and stored in 100% methanol at -20°C. On the day of staining, fish were rehydrated in decreasing concentrations of methanol (75%, 50%, 25%) and washed in phosphate buffered saline with 0.1% Tween (PBST). Larvae labeled with the presynaptic vesicle antibody SV2A (DSHB SV2-s, 52µg/mL, RRID: AB 2315387) were prepared as described elsewhere(*52*). Briefly, following rehydration, larvae were further permeabilized in PBS with 0.5% Triton and incubated in a blocking solution (1% BSA in PBT, 2.5% DMSO, and 5% HINGS). Larvae were then incubated for four nights in 3µg/µL of SV2A primary antibody at 4°C. Secondary antibody incubation was performed overnight at 1:1000 concentration in AlexaFluor 488nm goat anti-mouse (Invitrogen, A28175, RRID: AB 2535719). Alpha-bungarotoxin (*α*BTX) labeling was performed following a similar method as described previously(*53*). Following fixation and rehydration, larvae were incubated in 50µg /µL *α*BTX conjugated to AlexaFluor 647nm (ThermoFisher, B35450) for 45 minutes at room temperature in the dark. Fixed larvae double-labeled with SV2A and *α*BTX followed the SV2A antibody protocol as described above; following secondary antibody incubation and PBST wash, larvae were incubated in 50µg /µL BTX for 45 minutes prior to confocal imaging. Prior to labeling, all larvae were washed in PBST in the dark and mounted dorsal-side up in 2% low-melting point agarose (Thermo Fisher Scientific 16520) in E3 prior to imaging.

### Neuromuscular Junction Imaging and Quantification

Stacks of extraocular muscle were collected on a confocal microscope (Zeiss LSM800) with a 20x water-immersion objective (Zeiss W Plan-Apochromat 20x/1.0). The imaging window was centered on a single hemisphere over one eye at a time, 200–250 x 200–250 µm frame size. Stacks were collected through the full dorso-ventral extent of the muscle (*∼*30µm) with an imaging interval of 2µm. For all extraocular muscles, the imaging window was centered over one eye at a time with either both superior oblique and rectus (dorsal-up orientation) or both inferior oblique and rectus muscles (ventral-up orientation) in view.

Quantification of *α*BTX labeling in extraocular muscle was performed in FIJI/ImageJ (*49*). Maximum intensity projections of single-color grayscale stacks were made with the full muscle in view. Polygonal ROIs were drawn around the entirety of the extraocular muscle. Images were thresholded by intensity to differentiate the brightest signal (postsynaptic neuromuscular junction label) from the weakest signal (muscle fiber). Thresholding was repeated by two investigators independently to ensure consistency. Thresholded images were then binarized. “% Area Labeled” is defined as the fraction of suprathreshold pixels within each muscle’s ROI. Next, average particle sizes were computed with a set size of 1-infinity and converted to µm.

### Behavior Experiments & Analysis

Torsional eye rotations were measured in response to step tilts delivered using an apparatus similar in design to (*14*). All experiments took place in complete darkness. Larval fish were immobilized completely in 2% low-melting temperature agarose, and the left eye freed. The agarose was then pinned (0.1mm stainless minutien pins, FST) to a *∼*5mm^2^ piece of Sylgard 184 (Dow Corning) which was itself pinned to Sylgard 184 at the bottom of a 10mm^2^ optical glass cuvette (Azzota, via Amazon). Within age groups, body length was confirmed to be consistent to ensure matched developmental maturation (*54, 55*). The cuvette was filled with *∼*1mL of E3 and placed in a custom holder on a 5-axis (X,Y,Z,pitch,roll) manipulator (ThorLabs MT3 and GN2). The fish was aligned with the optical axes of two orthogonally placed cameras such that both the left utricle and two eyes were level with the horizon (front camera) and centered about the axis of rotation. The eye-monitoring camera (Guppy Pro 2 F-031, Allied Vision Technologies) used a 5x objective (Olympus MPLN, 0.1 NA) and custom image-forming optics to create a 100×100 pixel image of the left eye of the fish (6 µm/pixel), acquired at 200Hz. The image was processed on-line by custom pattern matching software to derive an estimate of torsional angle (LabView 2014, National Instruments), and data was analyzed using custom MATLAB scripts (Mathworks, Natick MA).

A stepper motor (Oriental Motors AR98MA-N5-3) was used to rotate the platform holding the cameras and fish. Microstep commands (0.0072°) were delivered with varying intervals to enable smooth speed control according to the algorithm in (*56*). An experiment consisted of 50 cycles of four steps each. Steps were *±*15° towards and away from the horizon. Each step followed a trapezoidal velocity profile peaking at either 35°/sec, peak acceleration 150°/sec^2^ (normal) or 90°/sec (fast controls), peak acceleration 600°/sec^2^. The platform velocity and acceleration were measured using integrated circuits (IDG500, Invensense and ADXL335, Analog Devices) mounted together on a breakout board (Sparkfun SEN-09268).

The eye’s response across the experiment was first centered to remove any offset introduced by the pattern-matching algorithm. Data were then interpolated with a cubic spline to correct for occasional transient slow-downs (i.e. missed frames). The eye’s velocity was estimated by differentiating the position trace; high-frequency noise was minimized using a 4-pole low-pass Butterworth filter (cutoff = 3Hz). Each step response was evaluated manually; trials with rapid deviations in eye position indicative of horizontal saccades or gross failure of the pattern-matching algorithm were excluded from analysis. The response to each step for a given fish was defined as the mean across all responses to that step across cycles. The gain was estimated by measuring the peak eye velocity occurring over the first second after the start of the step.

### Statistics

Parametric data were tested with Student’s t-test, paired t-tests for longitudinal data (i.e. comparing the same neuron), 1- and 2-way ANOVA with post-hoc comparisons, repeated measures design used for longitudinal data. Non-parametric data were tested using Kruskall-Wallis (K-W) tests. When appropriate, *α* was Bonferroni corrected for multiple comparisons, with a significance threshold of 0.05. A Kolmogorov-Smirnov (K-S) test was used to compare distributions of directionality indices. Population standard deviations were generated by bootstrapping (i.e resampling with replacement), n=60 repeats. Statistical tests were done on log-transformed data (dFF+1). All tests were repeated on raw data using non-parametric equivalents and the conclusions did not change, with one exception: in Figure 3J the 3–5 dpf differences between populations of neurons in *phox2a*^*-/-*^ and sibling fish changed from *p*_*ANOVA*_ = 0.053 to *p*_*K-W*_ = 0.0032. All tests reported in the text as subscripts; *p*_*post-hoc*_ refers to post-hoc pairwise comparisons after ANOVA/K-W using Tukey’s honestly significant difference procedure.

